# DOPAMINE D4 RECEPTOR DOWN-REGULATES RENAL SODIUM CHLORIDE COTRANSPORTER VIA UBIQUITINATION-ASSOCIATED LYSOSOME DEGRADATION

**DOI:** 10.1101/2024.02.14.580405

**Authors:** Mingzhuo Zhang, Mingda Liu, Zhiyun Ren, Weiwan Wang, Kiran K. Upadhyay, Laureano Asico, Ines Armando, Yutao Jia, Ping Wang, Ying Xue, Xiaoyan Wang

## Abstract

**Background:** The thiazide-sensitive sodium chloride cotransporter (NCC) is the major apical sodium transporter located in the mammalian renal distal convoluted tubule (DCT). The amount of sodium reabsorbed in the DCT through NCC plays an important role in the regulation of extracellular fluid volume and blood pressure. Dopamine and its receptors constitute a renal antihypertensive system in mammals. The disruption of *Drd4* in mice causes kidney-related hypertension. However, the pathogenesis of D4R-deficiency associated hypertension is not well documented.

**Method:** We assessed the effects of D4R on NCC protein abundances and activities of DCT in mice with renal or global *Drd4*-deficiencies and expressing human *D4.7* variant and in cultured mouse DCT cells, and explored the molecular mechanism.

**Results:** NCC inhibitor hydrochlorothiazide enhanced the natriuresis in *Drd4*^-/-^ mice. Renal NCC protein was greater while ubiquitination of NCC was less in *Drd4*^-/-^ than *Drd4*^+/+^ mice. Silencing of D4R in cultured mouse DCT cells increased NCC protein but decreased NCC ubiquitination. D4R agonist had opposite effects that were blocked by the antagonist. In mouse kidneys and DCT cells D4R and NCC colocalized and co-immunoprecipitated. Moreover, D4R-agonist promoted the binding between the two proteins demonstrated by fluorescence resonance energy transfer. D4R agonism internalized NCC, decreased NCC in the plasma membrane, increased NCC in lysosomes and reduced NCC-dependent-intracellular-sodium transport. The lysosomal inhibitor chloroquine prevented the D4R-induced NCC-reduction. A shortened NCC half-life was suggested by its decay under cycloheximide-chase. Ubiquitin-specific-protease 48 (USP48, a deubiquitinating enzyme) was increased in the kidneys and cells with *Drd4*-deficiency while D4R stimulation decreased it in *vitro* and reduction of USP48 with siRNA decreased NCC expression. The mice carrying human *D4.7* variant or with renal reduction of D4R developed hypertension with increased NCC.

**Conclusion:** Our data demonstrates that D4R downregulates NCC by promoting USP48-associated deubiquitination and subsequent internalization, lysosome relocation and degradation.

## INTRODUCTION

The thiazide-sensitive sodium chloride cotransporter (NCC) is the major apical sodium transporter located in the mammalian renal distal convoluted tubule (DCT)(1–3), encoded by *SLC12A3* gene and consists of the 1030 amino acid residues as a transmembrane protein. NCC is expressed both in the early DCT (DCT1) and in the late DCT (DCT2) in which the epithelial sodium channels (ENaC) with 3 subunits are also expressed(4, 5). Although the DCT accounts for only 7-10% of total sodium reabsorption in the mammalian nephron, mainly via NCC, the final urine composition and volume are normally determined in distal nephron segments, including the DCT. We have shown that NCC is a mediator of aldosterone escape, a phenomenon associated with pressure-natriuresis(6). Therefore, the amount of sodium reabsorbed in the DCT through NCC plays an important role in the regulation of extracellular fluid volume and blood pressure (BP). For example, decreased NCC function causes hypotension in Gitelman’s syndrome, whereas increased NCC function causes the hypertension in Gordon’s syndrome(7, 8). A NCC inhibitor, hydrochlorothiazide (HCTZ), is a common and effective diuretic(9) and recommended by American Heart Association and American College of Cardiology as one of first-line medicines for treatment of essential hypertension(10, 11).

Aldosterone targets NCC, ENaCs and water-channels to regulate the reabsorption of sodium, potassium, chloride and water from the distal nephron segments and finalizes the urine, through mineralocorticoid receptors(12, 13). Regulators of NCC include with-no-lysine-kinase 4 (WNK4)-oxidative stress-responsive gene 1-STE20-related proline/alanine-rich kinase pathways(14, 15) for NCC activation and phosphorylation, and Neural-precursor-cell-expressed-developmentally down-regulated protein 4-like (Nedd4-2) for NCC ubiquitination and subsequent degradation(16, 17). On the other hand, deubiquitylating enzymes may also affect the ubiquitination of sodium transporters sinceD3R-mediated inactivation of ubiquitin-specific-peptidase (USP48) prevents the removal of Ub-tags and promotes the degradation of the sodium hydrogen exchanger 3 (NHE3)(18), an apical sodium transport protein located in renal proximal tubules and thick ascending limbs of loop of Henle.

Dopamine and its receptors constitute a renal antihypertensive system in mammals(19, 20). A decrease in the renal synthesis of dopamine or dysfunction of any of the dopamine receptors causes hypertension(21, 22). The role of renal dopamine receptors in regulating sodium transport in the proximal tubule, thick ascending limb, and collecting duct is well established(22–26). All five dopamine receptor subtypes are present in the DCT(21). The increase of NCC is seen in the mice with gene deletion of *Drd2* and *Drd5*(27, 28). However, the effect of dopamine receptor D4R on sodium transporters in the DCT is not well established.

D4R is expressed in the DCT(29) where NCC is a major apical sodium transporter(6). The disruption of *Drd4* in mice causes kidney-related hypertension with normal serum/renal renin levels but persistent hypotensive effect to an angiotensin-II-type-I-receptor blocker; Thus, we have hypothesized that D4R may inhibit DCT sodium transport by decreasing the protein abundance and function of NCC and balance urinary sodium excretion. We tested this hypothesis in global *Drd4*-gene knockout mice (*Drd4*^-/-^), immortalized mouse renal DCT cells and in mice expressing the human *D4.7* variant, a model of decreased D4R function(30) or with renal *Drd4*-silencing.

## METHODS

### Sex as a biological variable

The status and regulation of NCC function and protein are different between sexes(31, 32), only male mice were used to exclude the involvement of sex difference in the current study.

### Mice with Drd4^-/-^, D4.7 and renal subcapsular Drd-siRNA, feeding, BP monitoring and HCTZ treatment

*Drd4* general knockout mice (6 months, 20^th^ generation in C57/BL6J background) were gifts from Dr. Pedro Jose (George Washington University, Washington, DC, USA). *D4.7* knock-in mice(33) (10 months, 15^th^ generations in C57/BL6J background) were gifts from Dr. Sergi Ferré (National Institutes of Health, Department of Health and Human Services, Baltimore, MD, USA). C57/BL6J mice (3 months) were grouped by simple-randomization and uninephrectomized 1 wk before renal subcapsular infusion via osmotic minipumps (100 μl capacity, 0.5μl/h for 7 days; Alzet; Durect Corp., Cupertino, CA, USA) filled with validated Drd4-siRNA or mock-siRNA (3μg/d) as reported previously(18). The mice were maintained in the animal facilities of with 12-h-dark and 12-h-light cycles.

Ration-feeding was used for the mice with normal salt (0.4% NaCl) diet providing5g food, 10 ml water per 25 g body weight in agar-gelled format each day(6, 28). In order to determine NCC activity in *vivo*, HCTZ, a specific inhibitor on NCC, was administered by intraperitoneal injection (30mg/kg, in PBS 150-180 μl/mouse) at 10AM(34). 6-h-urine samples were collected in metabolic cages for measurements of sodium and creatinine before and after HCTZ injection. The Cr concentrations in serum and urine were measured with Cr Assay Kit (Jiancheng, C011-2-1). The sodium, potassium, chloride concentrations were measured by an automatic biochemical analyzers (BS-2000M, Cobas 6000). Serum aldosterone levels were determined by ELISA (Jiancheng, H188).

BPs were measured in mice by telemetry, with a transmitter (model TA11PA-C20, DSI, St. Paul, MN) introduced in the left carotid artery, in an animal facility room (12 hrs on-off light cycle) specifically designed for such studies and were also recorded in anesthetized mice (35).

### Mouse DCT cells and treatments with Drd4-siRNA, Usp48-siRNA, D4R agonists and antagonists

Immortalized mouse DCT cells were gifts from Dr. Peter A. Freidman (University of Pittsburgh)(36). Previous studies have shown that these cells express NCC(37–40). The cells were characterized as DCT cells by positive NCC and D4R but negative NHE3, Na^+^-K^+^-2Cl^-^ cotransporter (NKCC2) and αENaC in current studies (**Figure S1**). The mycoplasma-free cells were cultured in Dulbecco’s Modified Eagle’s Medium (DMEM)-F-12 medium containing heat-inactivated 10% fetal bovine serum and 10 μg/ml of antibiotic-antimycotic mixture, at 37 °C in an incubator with humidified 5% CO2 and 95% air. The mouse DCT cells were seeded and grown in multi-well culture plates (70-80% confluence), were treated with D4R drugs(41) in DMEM-F-12 with 1% fetal bovine serum after serum-free culture for 1 h. Mouse *Drd4*-siRNA (F-5’-3’GAAGGGAGCGCAAGGCAAUTT, R-5’-3’AUUGCCUUGCGCUCCCUUCTT), Mouse *Usp48*-siRNA (F-5’-3’ GUUAAAGGCUGAUGAACCATT, R-5’-3’ UGGUUCAUCAGCCUUUAACTT) and mock siRNA (General Biosystems, China) were transfected into mouse DCT cells by Lipofectamine™ 3000 (Invitrogen, L3000015) (20 nM, time as indicated) in DMEM-F-12 with 10% fetal bovine serum. Total cell lysates (TCL) were prepared with protease inhibitors in the sucrose solution as described in the previous study(42). Cell viability was assessed using a cell counting kit (CCK-8, Beyotime, C0037) while cytotoxicity was determined by measuring the LDH (LDH assay, Beyotime, C0017) in the cell supernatants as previously described(42).

### Drugs and antibodies

The sources of drugs were: D4R agonist PD168077 (Tocris Bioscience, 1065/10) and antagonist L-745,870 (Tocris Bioscience, 1002/10)(43, 44); NCC inhibitor metolazone (Met) (MedChemExpress, HY-B0209); Hydrochlorothiazide (HCTZ) (Sigma, H2910); Cycloheximide (CHX) (MedChemExpress, HY-12320); Chloroquine (CQ) (MedChemExpress, HY-17589A). The rabbit polyclonal D4R and chicken polyclonal NCC antibodies were generated and affinity-purified in our laboratory. The specificity of D4R antibodies were reported previously with positive D4R in *Drd4*^+/+^ mouse brains but negative in *Drd4*^-/-^ ones by immunoblotting(45, 46). The polyclonal rabbit NCC antibody, was a gift from Dr. Mark Knepper (NHLBI); the same peptide sequence (PGEPRKVRPTLADLHSFLKQEG) was used as the immunogen in the generation of NCC antibody in chicken. The specificity of the NCC antibodies was also reported previously(27, 47) and characterized by apical membrane staining in mouse DCT and absence of binding in NCC-null mice (**Figure S2**) which were gift from Dr. Mark Knepper. The commercial primary antibodies were mouse monoclonal against α-sodium-potassium ATPase (αNKA) (Sigma, 05-369), ubiquitin (Cell Signaling, 3936) and GAPDH (Proteintech, 60004-1-Ig), β actin (Proteintech, 81115-1-RR), Nedd4-2 (Proteintech, 13690-1-AP), WNK4 (Proteintech, 22326-1-AP), USP48 (Bethyl Laboratories, A301-190A), Proteasome 26S subunit, ATPase, 6 (PSMC6) (Abcam, ab22639), Lysosomal membrane protein 1 (LAMP1) (Abcam, ab208943). The specificity of the antibody against USP48(18) was previously reported while the validation information of other commercial antibodies, with knockout, knockdown, or over-expression of the corresponding genes, was provided by the companies.

### Immunofluorescence and fluorescence resonance energy transfer (FRET) with confocal microscopy

Two pairs of mice left kidneys were fixed with 4% paraformaldehyde and embedded in the same paraffin block for co-immunostaining as reported previously(42, 48). After antigen retrieval with boiling in sodium citrate antigen repair solution, the sections were incubated with primary antibodies (negative controls without primary antibodies), then fluorescence-conjugated secondary antibodies. Nuclei were stained with DAPI or Hoechst. Similar staining procedures were applied on methanol-fixed cell slips. Images were taken with Zeiss confocal microscope (LSM 900, Carl Zeiss, Jena, Germany). In the cortex area, 4 views at 3, 6, 9, 12 o’clock positions were taken from each *Drd4*^+/+^ and *Drd4*^-/-^ mouse kidney. The immunofluorescence intensities were measured using image-J software. The similar co-immunostaining procedures were applied for FRET(48, 49). The energy transfer rate at 488 nm was calculated using Zeiss Zen 3.0 during bleaching at 555.

### Semi-quantitative immunoblotting

Whole kidney homogenates (WKH) and TCL were prepared with phosphatase/protease inhibitors as described previously(42). Plasma membrane-enriched fractions was achieved from TCL by centrifugation at 17,000 g for 30 min(47). Samples with equal amounts of proteins were separated by 10% SDS-polyacrylamide gel and transferred onto nitrocellulose membranes. The membranes were sequentially probed with the primary antibodies at 4°C overnight and corresponding horseradish peroxidase-coupled secondary antibodies for chemiluminescence detected using Tanon 5200 Muti system or IR-dye 680-(LI-COR Bioscience, C50317-02) or 800-labeled secondary antibodies (LI-COR Bioscience, C50449-01) for infrared signals detected and quantified with ODYSSEY Clx Licor and Image Studio Ver 5.2 system. A nice linear correlation was seen between loaded protein amounts and detected signals with both systems (**Figure S3**).

### Coimmunoprecipitation (co-IP)

Equal amounts of WKH or TCL (100 μg) were immunoprecipitated with protein G beads (Beyotime, coimmunoprecipitation kit, P2197S) with first primary (mouse or rabbit) antibodies and immunoblotted with second primary antibodies (different hosts)(42). Normal rabbit or mouse IgG as the immunoprecipitants were used as negative controls while TCL or WKH and target proteins were served as positive controls. The studies were performed at least three times.

### RNA extraction and real-time PCR

The procedures were similar as published(42). RNA was extracted from WKH and TCL. 1 μg RNA was used for reverse-transcription to get cDNA with HiScript III RT SuperMix (Vazyme R323-01). The quantitative PCR was performed on 1 μl cDNA for 40 cycles with ChamQ Universal SYBR qPCR Master Mix (Vazyme Q711) by Quant Studio 3 Applied Biosystems against the primers of *Drd4*, *Ncc* and *β-actin* (**Table S2**).

### Histology and immunohistochemistry with light microscopy

H&E and immunocytochemistry staining were performed as described(6). After dewaxing, hydration, and antigen retrieval, kidney sections were incubated sequentially with primary and secondary antibodies. At least 5 randomly chosen fields were viewed under Nikon Eclipse 80i microscope equipped with a digital camera (DS-Ri1, Nikon).

### Sodium transport study

Sodium transport was accessed, modified from previous reports(39, 40, 50), in DCT cells grown in Transwells on polyester membrane inserts (Corning Inc, #3460) by measuring intracellular sodium concentration with cell permeable Sodium Green^TM^ (Invitrogen, S6901)(51) instead of ^22^Na^+^(39, 40, 50). After the cells had achieved a polarized monolayer and pre-starved for 2 h, the serum-free medium was switched to an isotonic, chloride-free medium for 30 min to create a gradient for Na^+^ transport via the cotransporter. The vehicle or drugs in the chloride-free medium were added to the upper chambers (apical side) and allowed to incubate for 60 mins. Sodium Green^TM^ in chloride-containing medium along with ouabain (50 μM) was added to both upper and bottom chambers and incubated for 30 min at room temperature in the dark. The intracellular sodium concentration was quantified immediately with Victor V (fluorescence excitation at 485 nm, emission at 535 nm, 1420 Multilabel Counter, Perkin Elmer) and corrected for the protein concentration of the samples. A serial dilution of the cell suspension was generated in different wells to obtain a standard curve of intracellular sodium as a function of cell-count/protein-concentration. Initial sodium transport at 0, 5, 10, 15, 30, 60 min was determined.

### Biotinylation of cell membrane

After D4R agonist treatment, the cells on cover slips in cell culture plates were placed on ice and washed with ice-cold PBS, then incubated with cell-impermeable biotin (PIERCE, 21217) for 30 mins. 10 mM glycine in PBS was added. After washing, the cells were incubated with streptavidin-conjugated Alex 555 (red) for 15 min and fixed with cold 100% methanol for 20 min at -20℃. The cover slips were incubated with rabbit NCC primary antibody (1: 50 overnight in cold room), then secondary antibody (Alex 488, green) for 1 h. The rest procedures(48) were same as the co-immunostaining.

### Measurements of NCC half-life and degradation

Mouse DCT cells were grown to post-confluence in 6-well culture plates and treated with 50 μM CHX (a powerful protein synthesis inhibitor)(52). At each point time, cells were harvested with protease inhibitors. In additional cells, CQ, the lysosomal inhibitor, was added to the cultures in a final concentration of 20 μM in the presence or absence of PD168077 for 24 h. The rest procedures(6, 42) were similar as published for immunoblotting analysis.

### Statistical Analysis

Results were reported as mean ± SEM (standard error of the mean). Significant differences between two groups were determined by the student’s *t*-test. Significant differences among groups (> 2) were determined by one-way factorial ANOVA, followed by Holm-Sidak test. A value of *P* < 0.05 was considered statistically significant.

### Study approval

All procedures were conducted in accordance with NIH guidelines for the ethical treatment and handling of animals and ethics approval by the Animal Care and Use Committees of University of Maryland and Nanjing Medical University for research.

### Data availability

Relevant information about data available directly from the corresponding author.

## RESULTS

### Deletion of the *Drd4* gene in mice increased HCTZ-sensitive natriuresis without affecting serum aldosterone, creatinine clearance or serum creatinine

Sodium excretion 6-h after HCTZ injection was greater in *Drd4*^-/-^ mice than that in WT littermates (**Figure 1A**), suggesting that the *Drd4*^-/-^ mice had a higher HCTZ sensitivity than *Drd4*^+/+^ mice. SBPs were higher in *Drd4*^-/-^ mice than *Drd4*^+/+^ littermates at baseline, consistent with our previous results(35). There were no significant differences in the levels of serum creatinine, creatinine clearance rate (Ccr) or serum aldosterone between the mouse strains (**Figures 1B & 1C)**. The mRNA expression of *Ncc* tended to be higher in *Drd4*^-/-^ but was not significantly different (**Figure 1D**). However, the protein abundance of NCC, a major apical sodium transporter in DCT, was greater in homogenates of the cortex in *Drd4*^-/-^ than *Drd4*^+/+^ mice (left panel, **Figure 1E**). NCC and D4R co-stained in the apical membranes of the distal convoluted tubules of the kidneys (right panel, **Figure 1G**) and co-immunoprecipitated in whole kidney homogenates (**Figure 1F**). The physiological data of the mice including age, serum electrolytes, urine volume and electrolytes were similar in both mouse strains (**Table S1**). Negative controls for NCC and D4R immunostaining are shown in **Figure S4**.

**Figure 1.**
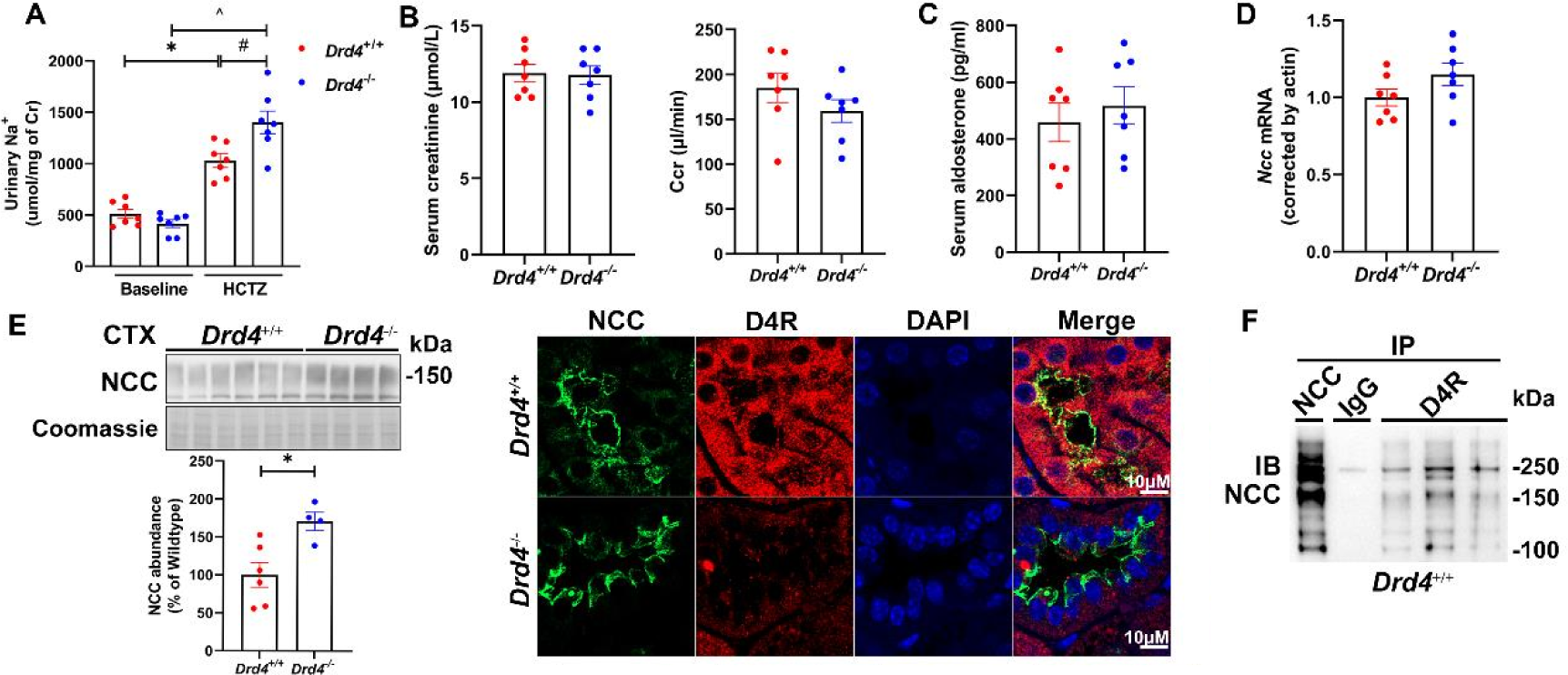
Sodium excretion in response to HCTZ, serum aldosterone and creatinine (Cr) from *Drd4*^-/-^ mice, colocalization between renal D4R and NCC. A. Hydrochlorothiazide (HCTZ) on urinary sodium excretion corrected by creatinine (Cr). The urinary sodium excretion in both mouse strains was increased relative to baseline while *Drd4*^-/-^ excreted more sodium than *Drd4*^+/+^ within 6-h of HCTZ injection (IP, 30 mg/kg). *^# *P* < 0.05, n = 7/group, one-way ANOVA, Holm-Sidak test. B. Serum creatinine levels and Cr clearance (Ccr) for glomerular filtration rates in *Drd4*^-/-^ and *Drd4*^+/+^ mice. The two groups had similar levels of serum Cr and Cr clearances. n = 7/group, Student’s *t*-test. C. Serum aldosterone levels were determined by ELISA from *Drd4*^-/-^ and *Drd4*^+/+^ mice. Aldosterone levels were similar in the two groups of mice. n= 7/group, Student’s *t*-test. D. *Ncc* mRNA levels measured by q-PCR in whole kidney homogenates from *Drd4*^+/+^ and *Drd4*^-/-^ mice. They were similar between the mouse strains. n = 7/group, Student’s *t*-test. E. Immunoblotting and co-immunofluorescent staining in the cortex (CTX) from in additional *Drd4*^-/-^ (n=4) and *Drd4*^+/+^ (n=6) mice. NCC was higher in *Drd4*^-/-^ than *Drd4*^+/+^ group. \**P* < E. 0.05, Student’s *t*-test (left panel). Under laser confocal microscopy, D4R (red, rabbit antibody, 1:50) was not shown in NCC-positive (green, chicken antibody, 1:50) renal tubules of *Drd4*^-/-^ mice but was expressed diffusely in renal tubules in *Drd4*^+/+^ mice. NCC (distal convoluted tubule marker) and D4R colocalized (yellow) in the apical membranes of the distal convoluted tubules. The nuclei were stained by DAPI. 400x, scale = 10μM (right panel). F. Co-immunoprecipitation of D4R and NCC. Equal amounts (100 μg) of whole kidney homogenates from *Drd4*^+/+^ mice were immunoprecipitated (IP) with anti-D4R antibodies (rabbits, 1 μg) while normal rabbit IgG (1 μg) was used as IP negative control and NCC (rabbits, 0.5 μg) antibody was used as IP positive control. Immunoblotting was performed against NCC (Chicken, 1:1000) antibody. NCC was seen in D4R pull-downs from 3 different mouse kidneys.

### Silencing of the *Drd4* gene in cultured mouse DCT cells increased NCC and USP48 expression and decreased NCC ubiquitination

We used immortalized mouse renal DCT cells to elucidate the mechanism leading to decreased NCC in the presence of lack D4R function. Treatment with *Drd4*-siRNA (20 nM) for 48 h reduced the protein abundance of D4R by about 40% indicating a moderate gene silencing efficiency. The protein abundance of factors that may be involved in the regulation of NCC expression were studied in the DCT cells. αNKA (an abundant basolateral sodium pump in DCT), and Nedd4-2 were not altered but USP48 was higher (**Figures 2A & 2B**) and ubiquitinated-NCC level was lower in *Drd4*-siRNA than in the mock treated cells (**Figures 2C & 2D**). At mRNA level *Drd4* was lower while *Ncc* was higher (**Figure 2E**) in *Drd4*-siRNA-treated than in mock-treated cells. The sequences of *Drd4*-siRNA, the detailed information on the primers (**Table S2**) and melting curves of qPCR products (**Figure S5**) are included in the supplemental file.

**Figure 2.**
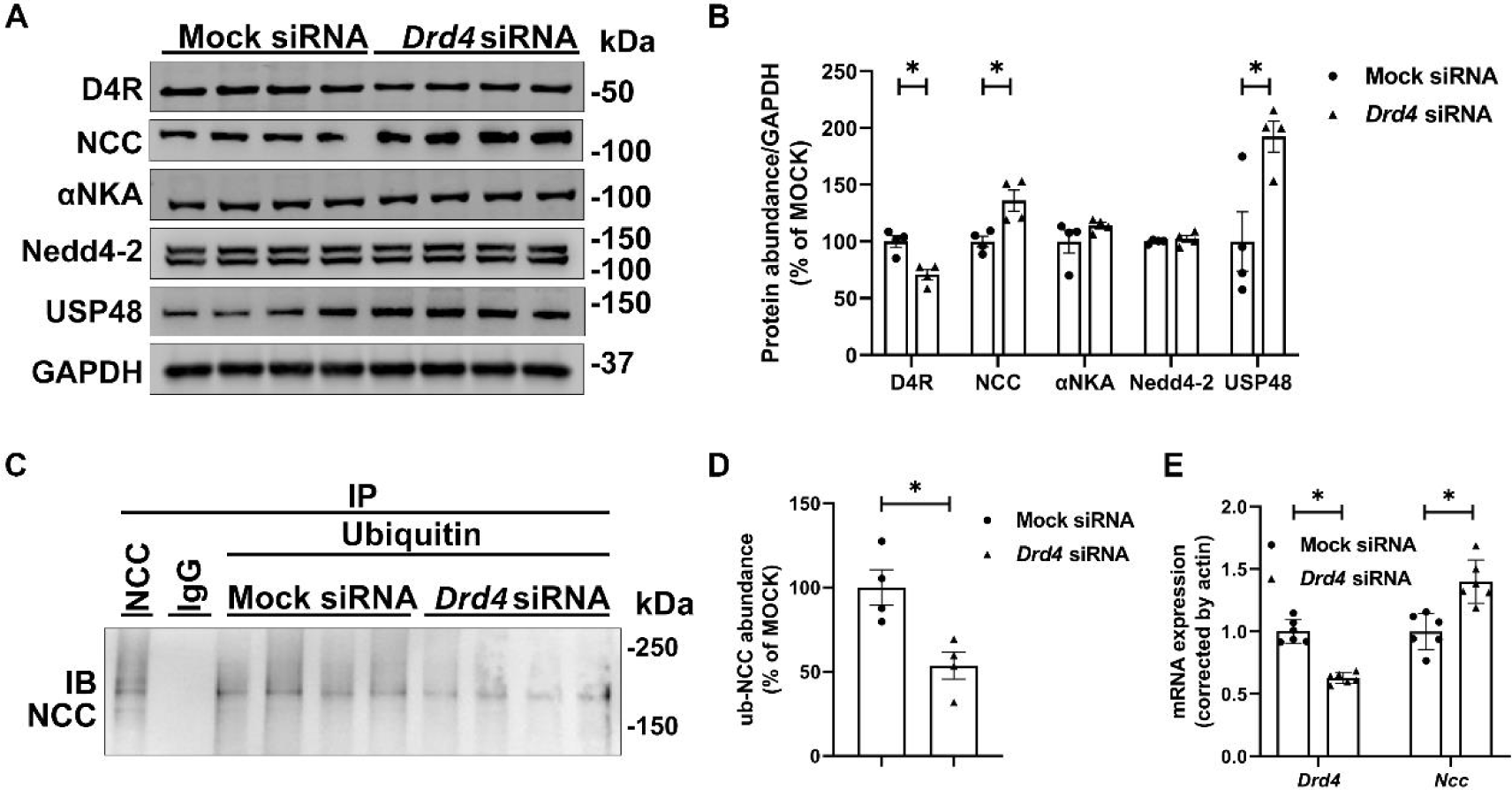
NCC, NCC ubiquitination and *Ncc* mRNA in mouse renal DCT cells treated with *Drd4*-siRNA for 48 h. A. Immunoblotting of D4R, NCC, αNKA, Nedd4-2, USP48 and GAPDH in total cell lysates (TCL) from mouse DCT cells treated with mock or *Drd4*-siRNA (20 nM, 48 h). B. Quantitative analyses of the immunoblots corrected by GAPDH. D4R was lower but NCC and USP48 were higher in *Drd4*-siRNA than mock group. αNKA and Nedd4-2 were not altered. \**P* < 0.05, n = 4/group, Student’s *t*-test. C & D. Co-immnoprecipitation between ubiquitin and NCC in TCL of mouse DCT cells treated with mock or *Drd4*-siRNA using mouse IgG as IP negative control and NCC antibody as IP positive control. ubiquitinated NCC (ub-NCC) was lower in *Drd4*-siRNA than mock group. \**P* < 0.05, n = 4/group, Student’s *t*-test. E. mRNA levels of *Drd4* and *Ncc* measured by QPCR. *Drd4* mRNA was decreased while *Ncc* mRNA was increased by *Drd4*-siRNA treatment, \**P* < 0.05, n = 6/group, Student’s *t*-test.

### D4R agonism with PD168077 decreased NCC and USP48 expression and promoted NCC ubiquitination in mouse DCT cells

The D4R agonist PD168077 lowered NCC protein abundance in mouse DCT cells but did not affect αNKA, in a concentration-dependent manner (**Figures 3A**). The cell viability and toxicity were not affected by PD168077 at concentrations of 0, 5,10 μM, however, at 20, 40 μM the drug decreased cell growth and increased LDH release rates (**Figure S6**). NCC and D4R coimmunoprecipitated in mouse DCT cells and the coimmunoprecipitation was increased at 10 μM but not at 1 μM PD168077 (**Figure 3B**). There was a small but significant decrease in USP48 while Nedd4-2 was not affected by PD168077 at 10 μM (**Figures 3C**). The agonist promoted the ubiquitination of NCC in mouse DCT cells (**Figures 3D**). PD168077-induced NCC decrease was blocked by the D4R antagonist L-745,870 (**Figure 3E**).

**Figure 3.**
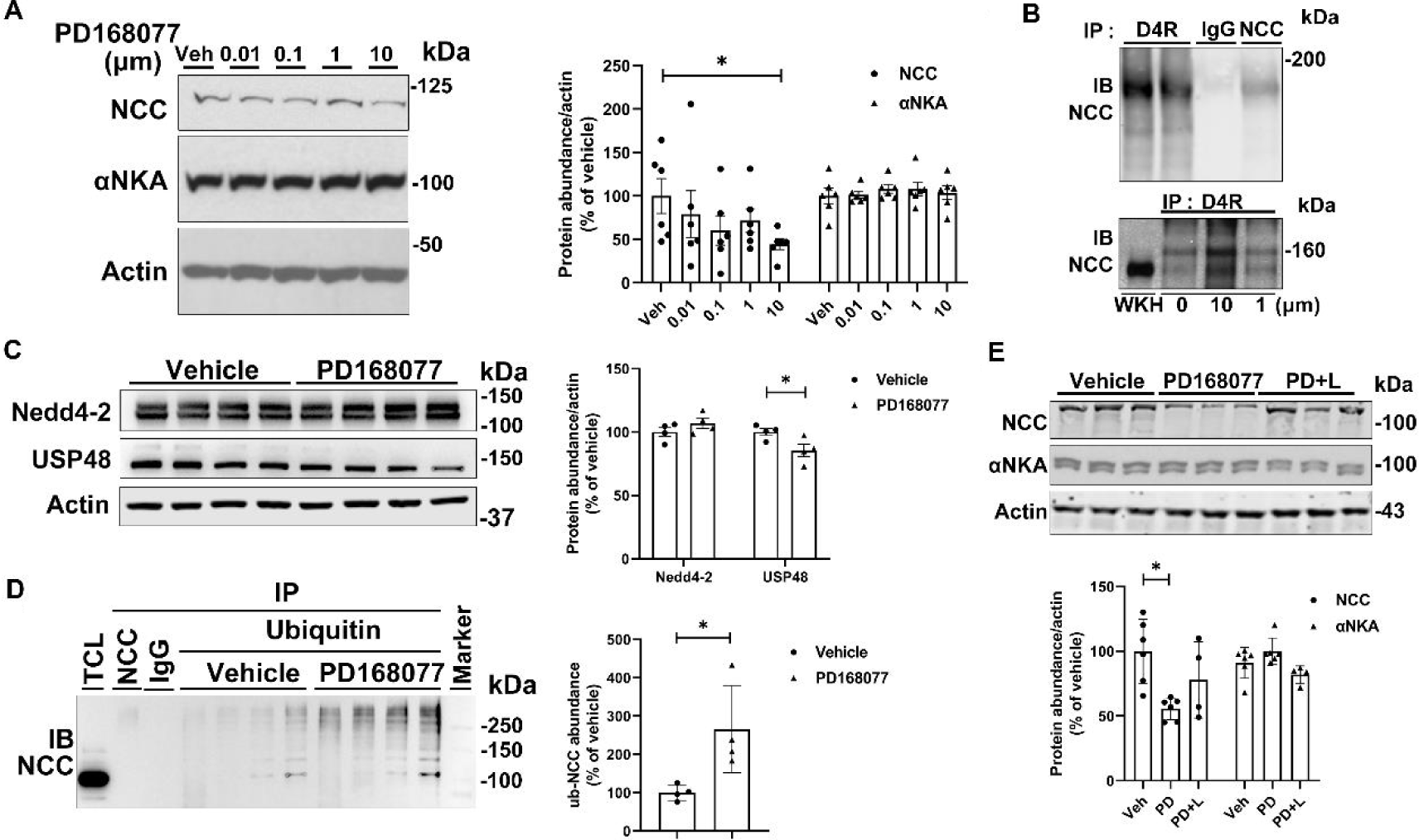
NCC and its ubiquitination, coimmunoprecipitation between D4R and NCC in mouse renal DCT cells treated with D4R agonist for 24 h. A. Immunoblotting and quantitative analyses of NCC and αNKA in TCL from mouse renal DCT cells treated with a D4R agonist PD168077 (PD) at 0.01, 0.1, 1 and 10 μM. The protein abundance of NCC, not αNKA, (corrected by actin) tended to decrease by the D4R agonist and reached the significance at 10 μM. \**P* < 0.05 vs vehicle (Veh), one-way ANOVA, Holm-Sidak test, n = 6/concentration. B. Coimmunoprecipitation of D4R and NCC. Upper panel: Equal amounts of TCL proteins from mouse DCT cells were immunoprecipitated (IP) with two different in-house anti-D4R antibodies (rabbits 9932, 9950, 1: 50) while normal rabbit IgG (1 μg) was used as IP negative control and NCC (0.5 μg) antibody was used as positive control. D4R co-immunoprecipitated with NCC in vehicle-treated mouse DCT cells. Lower panel: TCL of mouse DCT cells, pre-treated with vehicle or PD (10 μM and 1 μM, 24 h), were immunoprecipitated with anti-D4R antibody. Whole kidney homogenate (WKH) from *Drd4*^+/+^ mice was served as positive IB control. PD, at 10 μM but not 1 μM, increased the coimmunoprecipitation of D4R and NCC, relative to vehicle (0 μM). C. Immunoblotting and quantitative analyses of Nedd4-2 and USP48 in mouse DCT cells treated with vehicle or PD (10 μM, 24 h). USP48, not Nedd4-2, was reduced by the D4R agonist. D. U-NCC. Co-immnoprecipitation between ubiquitin and NCC in TCL of mouse DCT cells treated with vehicle or PD using mouse IgG as IP negative control, NCC antibody as IP positive control and TCL as IB positive control. ub-NCC was higher in PD than vehicle treated group. \**P*<0.05, n = 4/group, Student’s *t*-test. E. Immunoblotting and quantitative analyses of NCC in mouse DCT cells treated with D4R agonist PD and antagonist L-745,870 (L) for 24 h. The protein abundance of NCC is decreased by PD (10 μM) but the decrease is prevented by the co-incubation of the D4R agonist with the L (10 μM). The drugs did not affect the abundance of αNKA. One-way ANOVA, Holm-Sidak test, n = 4-6/treatment, \**P* < 0.05 vs. Others.

### D4R agonism internalized NCC, relocalized it to lysosomes and decreased apical sodium transport

Within 1 h incubation, the D4R agonist PD168077 moved NCC staining from the plasma membrane into the cytosol (**Figure 4A**) and decreased NCC but not αNKA protein in plasma membrane-enriched (PM) fractions **(Figures 4B & 4C**). Initial Cl^-^ dependent Na^+^ transport was detected in the upper chamber (apical side) of polarized mouse DCT cells grown in Transwells for intracellular sodium accumulation (**Figure S7**). Sodium transport at 0, 5, 10, 15, 30, 60 min (R=0.83, *P*<0.05) in untreated DCT cells followed a linear pattern (**Figure 4D**). PD168077 decreased apical sodium transport while D4R antagonist L-745,870 blocked, partially at 1 μM and fully at 10 μM, the D4R-mediated inhibition of apical sodium transport (**Figure 4E**), similarly but not synergistic to that caused by metolazone (Met), a thiazide-like diuretic (**Figure 4F**). Treatment with PD168077 for 1 h while decreased NCC staining in the plasma membrane increased its signal in the cytoplasm without changes in D4R staining. The color of their colocalization from yellow turned to orange in the merged image due to the reduction of green (NCC) signal (**Figure 5A**). PD168077 induced changes in NCC location were blocked by D4R antagonist (**Figure 5B**). After 1 h of treatment NCC was internalized from biotinylated plasma membrane relative to baseline (**Figure 5C**). The co-staining between NCC and proteasomal marker proteasome 26S subunit, ATPase 6 (PSMC6) was minimal and unchanged with PD168077 for 1h (**Figure 5D**). D4 agonism also promoted NCC to relocate into lysosomal membrane protein 1 (LAMP1) in an area close to the plasma membrane after 1 h of incubation with PD168077 (**Figure 5E**). The energy transfer rate between NCC and D4R with FRET showed that PD168077 enhanced the FRET efficiency after 1 h treatment relative to 0 min (**Figure 5F**). The representative images for the colocalization and energy transfer curves were included in **Figure S8**.

**Figure 4.**
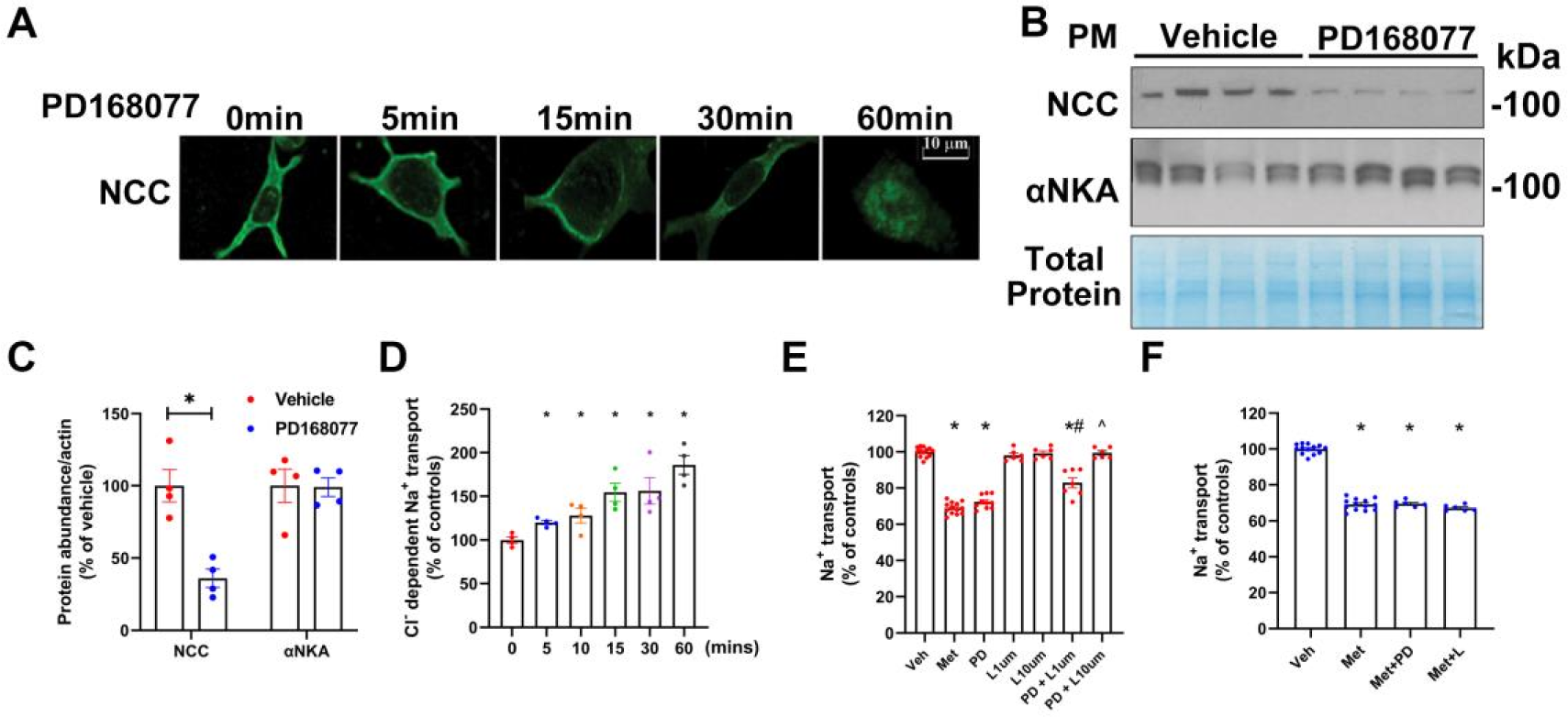
NCC protein and activity in mouse DCT cells treated with D4R agonist PD168077 (10 μM) within 60 min. A. Time course of NCC under confocal microscopy on mouse DCT cells treated with PD from 0-60 min as indicated. NCC (green, chicken, 1:100) is mainly located in the plasma membrane with some staining in the cytoplasm at baseline. The D4R agonist appeared to decrease NCC staining in the plasma membrane at 30 & 60 min while the staining in the cytoplasm was diffused with the time. B & C. Immunoblotting and quantitative analyses of NCC and αNKA protein in plasma membrane (PM)-enriched fraction of mouse DCT cells treated with a D_4_R agonist for 60 min. The protein abundance of NCC, not αNKA, (corrected by actin) was decrease by the D4R agonist. Equal loading was confirmed by total protein staining with Commassie blue. \**P* < 0.05, Student’s *t*-test, n = 4/group. D. Cl-dependent Na^+^ transport. Initial apical Na+ transport (upper chamber) of mouse DCT cells in presence of ouabain measured by the fluorescence of sodium green at different time points. Na^+^ transport into the cells rose with time (R=0.83, *P*<0.05). E. Apical sodium transport in polarized mouse DCT cells treated with D4R agonist PD for 60 min. PD (10 μM) decreased intracellular sodium, similar to the NCC inhibitor Met. The D4R antagonist L-745,870 (L, 10 μM) did not inhibit sodium transport by itself (96.6±0.5%, n = 4), but decreased (PD + L) the agonist (PD)-mediated inhibition partially at 1 μM and completely at 10 μM. All drugs were added in the upper chambers of the Transwells. *P* < 0.05 vs. Veh*, Met^#^ or PD^ alone, one-way ANOVA, Holm-Sidak test. F. The apical sodium transport in polarized mouse DCT cells treated with D4R agonist and antagonist combined with Met. Neither D4R agonist nor D4R antagonist at 10 μM for 60 min affected the met - mediated inhibition of sodium transport. *P* < 0.05 vs. vehicle*, one-way ANOVA, Holm-Sidak test, n = 6-9/group.

**Figure 5.**
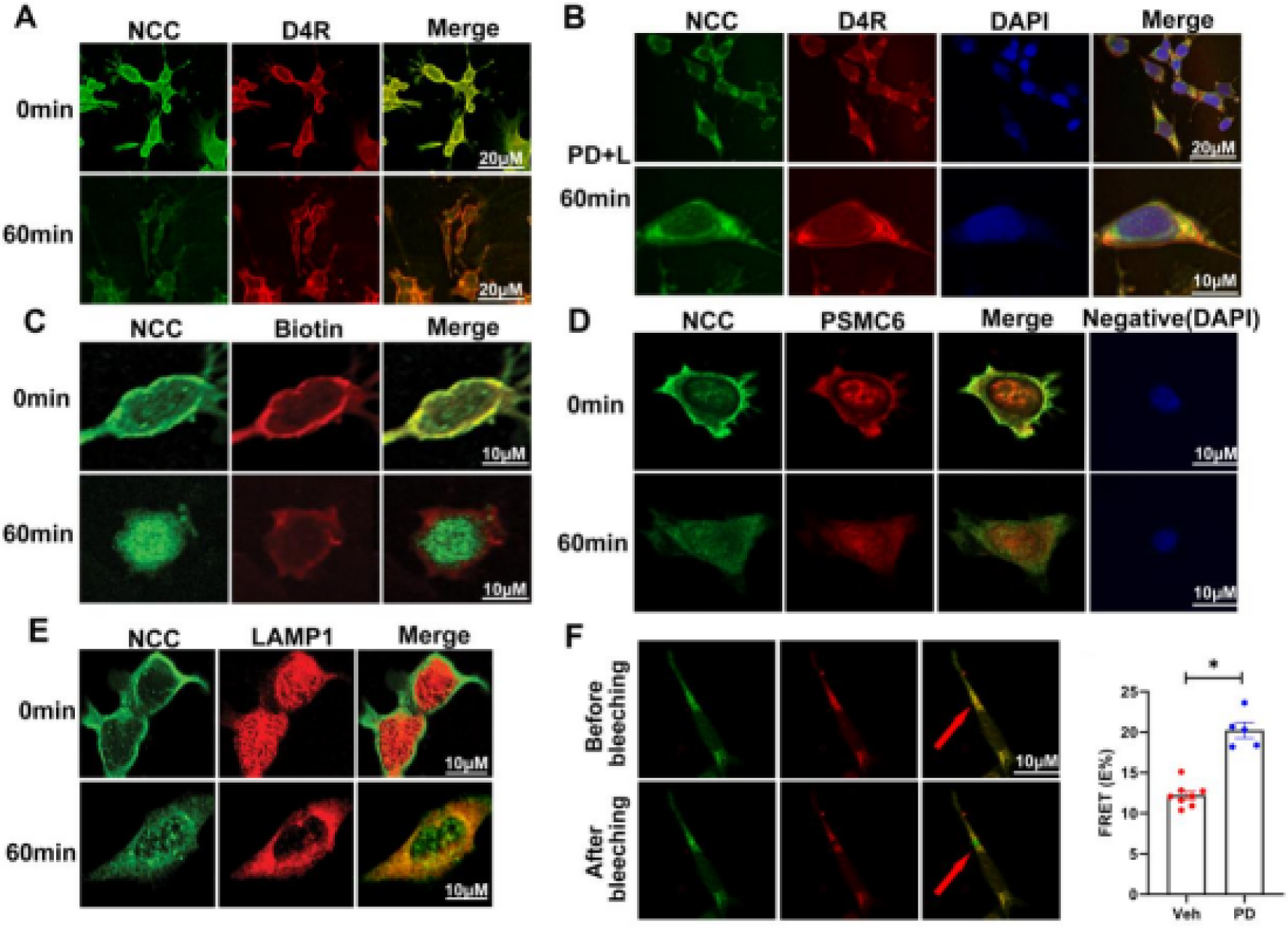
NCC internalization and colocalization with D4R, Lysosomal membrane protein 1 (LAMP1), proteasome 26S subunit, ATPase 6 (PSMC6) in mouse DCT cells treated with D4R agonist PD168077 for 60 min. A. Co-staining of NCC with D4R in mouse DCT cells treated with PD168077 (10 μM, 60 min). NCC staining (green, chicken, 1:500) in the plasma membrane appeared to be decreased while the staining in the cytoplasm appeared to be increased by the treatment. D4R staining (red, rabbit, 1:100) was not altered by the agonist, still located in the plasma membrane with some staining in the cytoplasm. The color of their colocalization from yellow turned to orange in the merged image due to the reduction of green (NCC) signal. Scale = 20 μM. B. The co-incubation of the agonist PD168077 (PD, 10 μM) and antagonist L-745,870 (L, 10 μM) for 60 min. After the co-incubation both NCC (green) and D4R (red) remained in the plasma membrane and cytosol but their colocalization was mainly close to the plasma membrane. Relative to D4R, more NCC remained in the perinucleus. The scale was 10 μM (top panel) and 20 μM (bottom panel). C. Internalization of NCC. NCC (green, chicken, 1:500) was co-stained with the biotinylated plasma membrane (red) at 0 min. However, the costaining was abolished by the D4R agonist PD 168077 (10 μM, 60min), associated with internalization of NCC (green) relative to the biotinylated plasma membrane (red). Scale = 10 μM. D. Colocalization of NCC with proteasome marker (PSMC6). At 0 min, there was minimum colocalization of NCC (1:500, chicken, green) and PSMC (1:50, mouse, red) in the plasma membrane and cytosol. PD168077 appeared to decrease NCC and PSMC6 at the membrane but did not increase their colocalization (bottom panel). Negative control was stained without the primary antibodies under same conditions (secondary antibodies only). Nucleus was stained with DAPI. The scale was 10 μM. E. Colocalization of NCC with lysosome marker LAMP1. At 0 min, there was no colocalization of NCC with LAMP1 since NCC (1:500, chicken, green) was located mainly in the plasma membrane and LAMP1 (1:50, mouse, red) was in cytosol (top panel). After 60 min of incubation with the D4R agonist, PD168077 (PD, 10 μM), NCC immunostaining decreased at the plasma membrane but increased in the cytosol and became colocalized (yellow in the merge image) with LAMP1 (bottom panel). Scale = 10 μM. F. The energy transfer rate technique (FRET) for the colocalization (yellow) of NCC (green) and D4R (red) before and after photo bleaching at wavelength 555. PD168077 (n=5) enhanced the FRET efficiency between NCC and D4R after 60 min treatment relative to 0 min (n=8). \**P* < 0.05, Student’s *t*-test.

### D4R agonism decreased NCC half-life and promotes its lysosomal degradation in mouse DCT cells

NCC protein abundance was reduced along with cycloheximide (CHX)-chase at 2, 4, 6,8h in the presence of PD168077 in a time-dependent manner (**Figures 6A & 6B**). The area under the curve for NCC decay with CHX-chase was reduced by PD168077 within8h (**Figure 6C**), indicating a shorten NCC half-life due to an increased protein degradation. PD168077-induced NCC decrease was reversed by CQ, a lysosomal inhibitor (**Figures 6D & 6E**), demonstrating that D4R-mediated NCC protein degradation may be lysosome-dependent. *Usp48*-siRNA mediated silencing for 48 h was concentration dependent and maximal at 20 nM.

**Figure 6.**
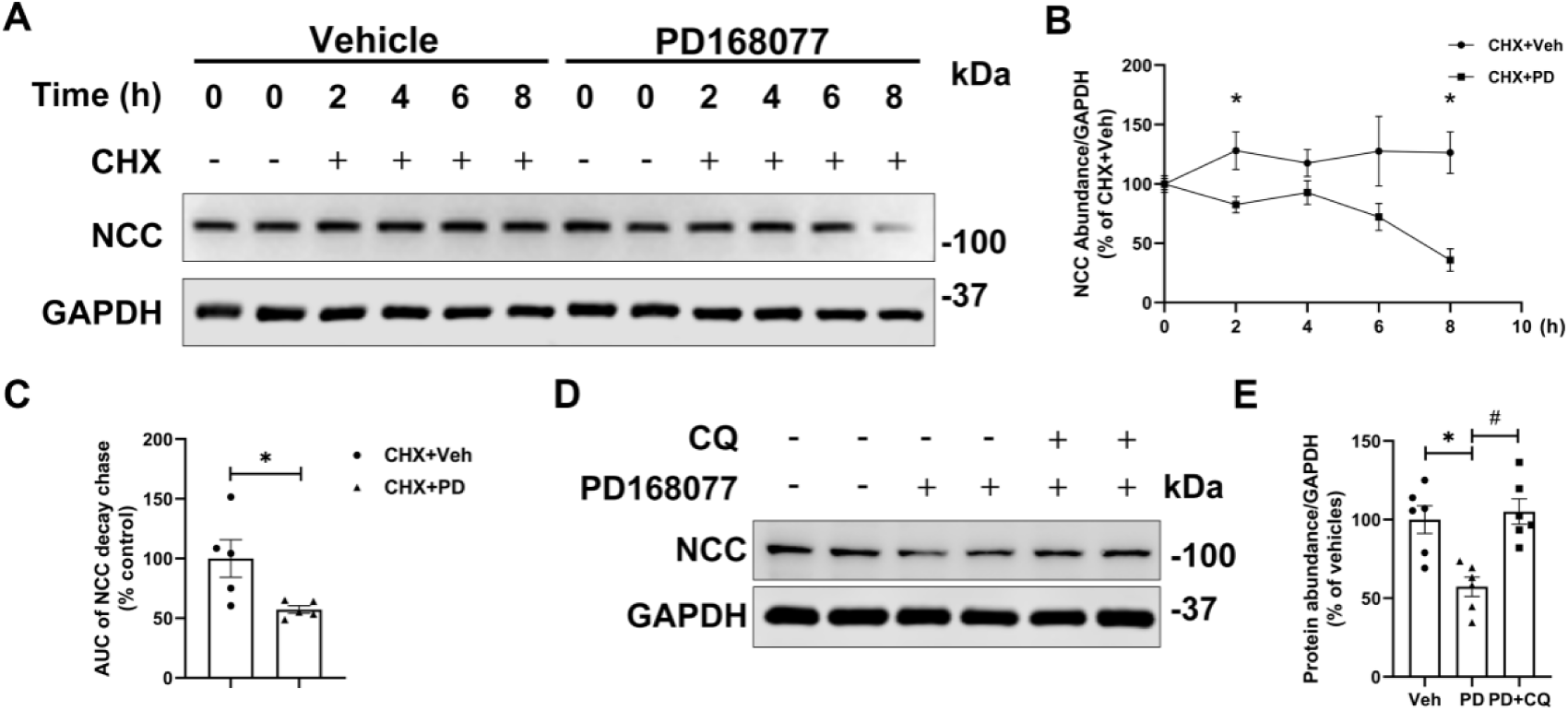
NCC degradation in mouse renal DCT cells treated with D4R agonist PD168077. A & B. Immunoblotting and quantitative analyses of NCC, corrected by GAPDH, with 8 h-CHX chase. NCC protein abundances tended to decrease by D4R agonist time-dependently in the presence of CHX. The significant differences were observed at 2, 8 h. \**P* < 0.05, n = 5, Student *t*-test. C. Area under the curve (AUC) for NCC degradation by PD168077 with 8-h CHX chase. PD168077 reduced NCC half-life by 50% relative to vehicle. \**P* < 0.05, n = 5, Student *t-*test. D & E. Immunoblotting and quantitative analyses of NCC, corrected by GAPDH, in mouse DCT cells treated with PD168077 and CQ for 24 h. CQ, a lysosomal inhibitor, prevented the PD168077-mediated NCC inhibition. \**P* < 0.05, one-way ANOVA, Holm-Sidak test, n = 6.

### *Drd4*^-/-^ mice exhibited increased USP48 expression and NCC ubiquitination

The protein abundance of NCC, a major apical sodium transporter in DCT(53), was greater in *Drd4*^-/-^ than *Drd4*^+/+^ mice while αNKA expression was not altered. USP48 was also higher in *Drd4*^-/-^ than *Drd4*^+/+^ mice while Nedd4-2(54) (a major ubiquitinating enzyme for NCC) and WNK4(55) (a major phosphorylation enzyme for NCC) were similar between strains (**Figures 7A & 7B**). NCC appeared to be concentrated in the apical membrane of DCT (**Figure 7C**). Low magnification images were provided in **Figure S9**. The immunofluorescent staining of NCC in the renal cortex was stronger in *Drd4*^-/-^ than *Drd4*^+/+^ mice (**Figures 7D**). The ubiquitinated NCC was higher in *Drd4*^-/-^ than *Drd4*^+/+^ mice (**Figures 7E**).

**Figure 7.**
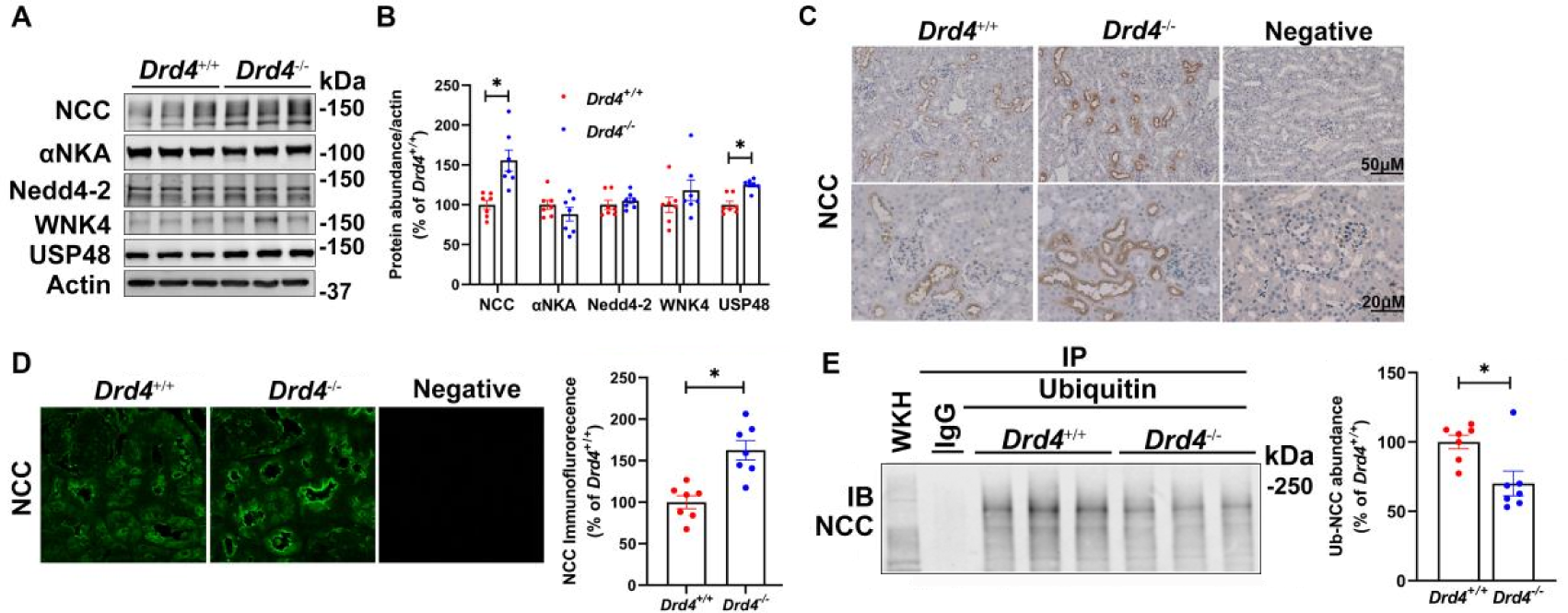
NCC, regulators, post-translational modifications in kidneys from *Drd4*^-/-^ and *Drd4*^+/+^ mice. A&B. Immunoblotting and quantitative analyses of αNKA, Nedd4-2, USP48, WNK4, and actin in whole kidney homogenates from *Drd4*^-/-^ and *Drd4*^+/+^ mice. The protein abundances of USP48, not αNKA, Nedd4-2 and WNK4 were higher in *Drd4*^-/-^ than *Drd4*^+/+^ mice. \**P* < 0.05, vs *Drd4*^+/+^, n = 7/group, Student’s *t*-test. C. Immunocytochemistry staining of NCC in the distal convoluted tubules of *Drd4*^+/+^ and *Drd4*^-/-^ mice under light microscopy. The brown staining of NCC in the apical membrane appeared more concentrated in *Drd4*^-/-^ than *Drd4*^+/+^ mice. 400x, scale = 20μM, 200x, scale = 50μM. D. Immunofluorescence of NCC in the renal cortex of *Drd4*^+/+^ and *Drd4*^-/-^ mice under confocal microscopy. *Drd4*^-/-^ showed stronger immunofluorescent staining of NCC (green, chicken antibody, 1:50) than *Drd4*^+/+^ mice. No signals were detected in the negative control under the same conditions without primary antibody. 400x, scale = 50μM. E. Quantitative analyses of the protein abundances of ub-NCC in whole kidney homogenates from *Drd4*^+/+^ and *Drd4*^-/-^ mice. Co-immnoprecipitation (IP) of ubiquitin (ub) with NCC (immunoblotting, IB) was performed with mouse IgG as negative IP control and WKH as positive IB control. 100 μg of protein was used from each mouse kidney homogenate. The protein abundances of ub-NCC was higher in *Drd4*^+/+^ than *Drd4*^-/-^ mice. \**P* < 0.05, n = 7/group, Student’s *t*-test.

### USP48 coimmunoprecipitated and co-stained with NCC in distal convoluted tubules of the mouse kidney

In wild-type littermates, USP48 and NCC co-immunoprecipitated in the whole kidney homogenates (**Figure 8A**) and co-stained in the distal convoluted tubules in mouse kidneys (**Figure 8B**). In mouse DCT cells treatment with *USP48*-siRNA (10, 20, 40, 80 nM) for 48 h decreased the levels of USP48 maximally at 20 nM while the expression of NCC was maximally decreased from 10 to 80 nM indicating that ubiquitination is necessary for NCC degradation (**Figures 8C & 8E**).

**Figure 8.**
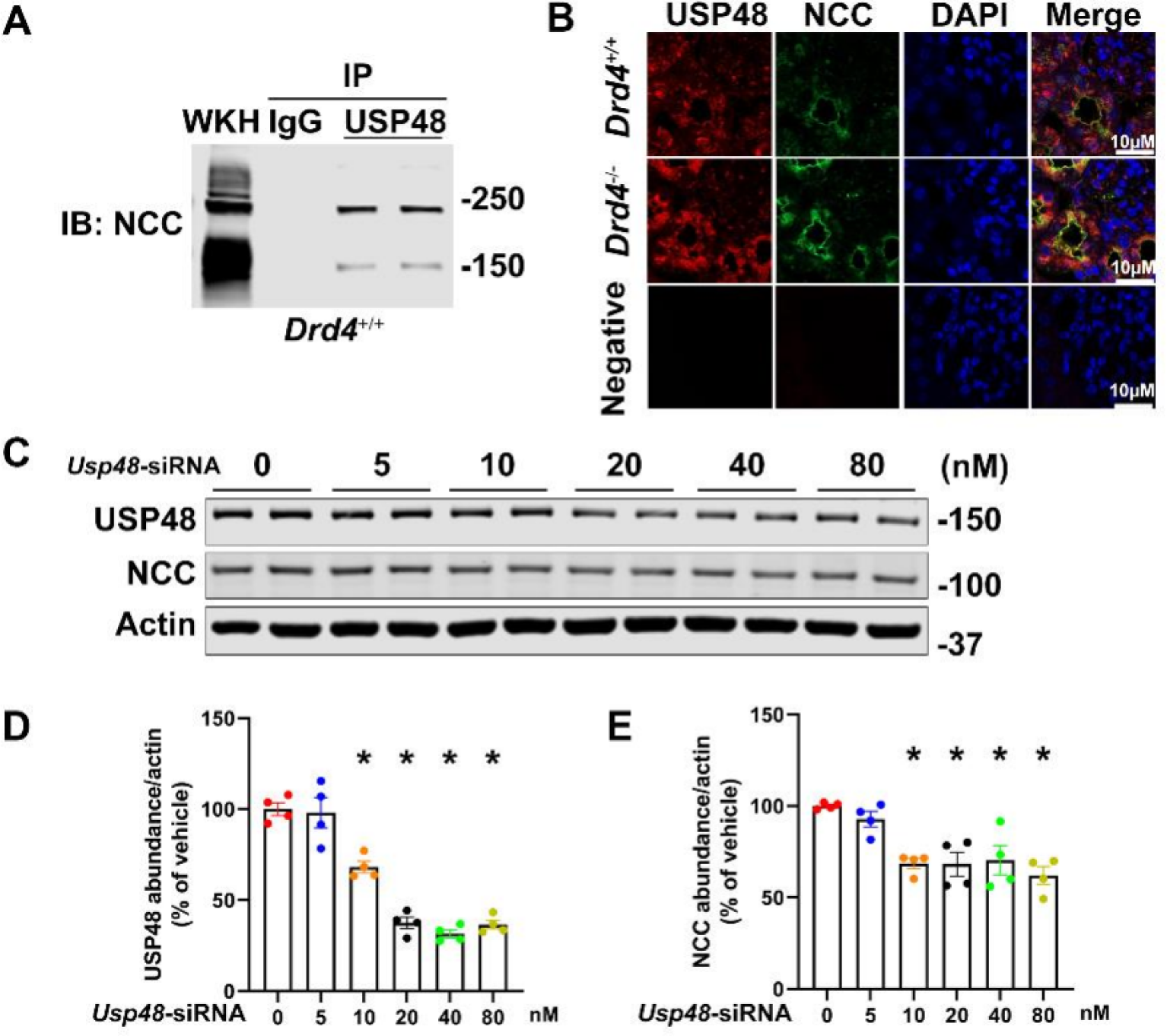
NCC and USP48 in mouse kidneys and DCT cells. A. Equal amounts (100 μg) of whole kidney homogenates from *Drd4*^+/+^ mice were immunoprecipitated (IP) with anti-USP48 antibodies (rabbits, 1 μg) while normal rabbit IgG (1 μg) was used as IP negative control. Immunoblotting was performed against NCC (Chicken, 1:1000) antibody. B. Immunofluorescence of USP48 and NCC under confocal microscopy in the renal cortex of *Drd4*^-/-^ and *Drd4*^+/+^ mice. USP48 (red, rabbit antibody, 1:50) and NCC (green, chicken antibody, 1:50) were colocalized (yellow) in the distal convoluted tubules. The nuclei were stained by DAPI. 400x, scale = 10μM. C-E. Immunoblotting and quantitative analyses of NCC, corrected by actin, in mouse DCT cells treated with different concentrations of *Usp48-*siRNA for 48h. USP48 protein abundances were declined by siRNA in the mouse DCT cell concentration-dependently (R=0.9022, P<0.05 by Pearson analysis) until 40 nM relative to 0 (mock-siRNA) but there was no further inhibition at 80 nM. However, the concentration dependent inhibition was not seen with NCC that was decreased similarly from 10 to 80 nM of siRNA. \**P* < 0.05 vs. Mock-siRNA (0 nM), n = 4, one-way ANOVA, Holm-Sidak test.

### The human D4R variant *D4.7* expressed in mice or kidney-specific-*Drd4* inhibition increased NCC protein abundance in the kidney and blood pressure

The human *D4R-gene* is highly polymorphic. The main polymorphisms are a 2 to 11 tandem repeats in a location coding for the third intracellular loop of the receptor. The variants containing two, four and seven repeats (*D4.2*, *D4.4* and *D4.7*) are the most frequently expressed(56). Knock-in mice carrying *D4.7* gene were shown to have behavior abnormalities(57). Stimulation of the *D4.7* variant appears to be less potent than the D4R wild type in inhibiting cAMP(33, 58). Systolic BP measured by telemetry in knock-in mice carrying the *D4.7* gene showed a persistent increase in the mutant mice at all times measured although urinary Na^+^ excretion was not different between the groups (**Figures 9A & 9B**). However, the renal expression of NCC in *D4.7 mice* was significantly higher than in mice expressing the wild type *D4* gene (**Figures 9C & 9D**). The renal reduction of D4R (30%) by subcapsular siRNA infusion increased BP and renal NCC in mice without changes in urine sodium excretion (**Figures 9E & 9F**).

**Figure 9.**
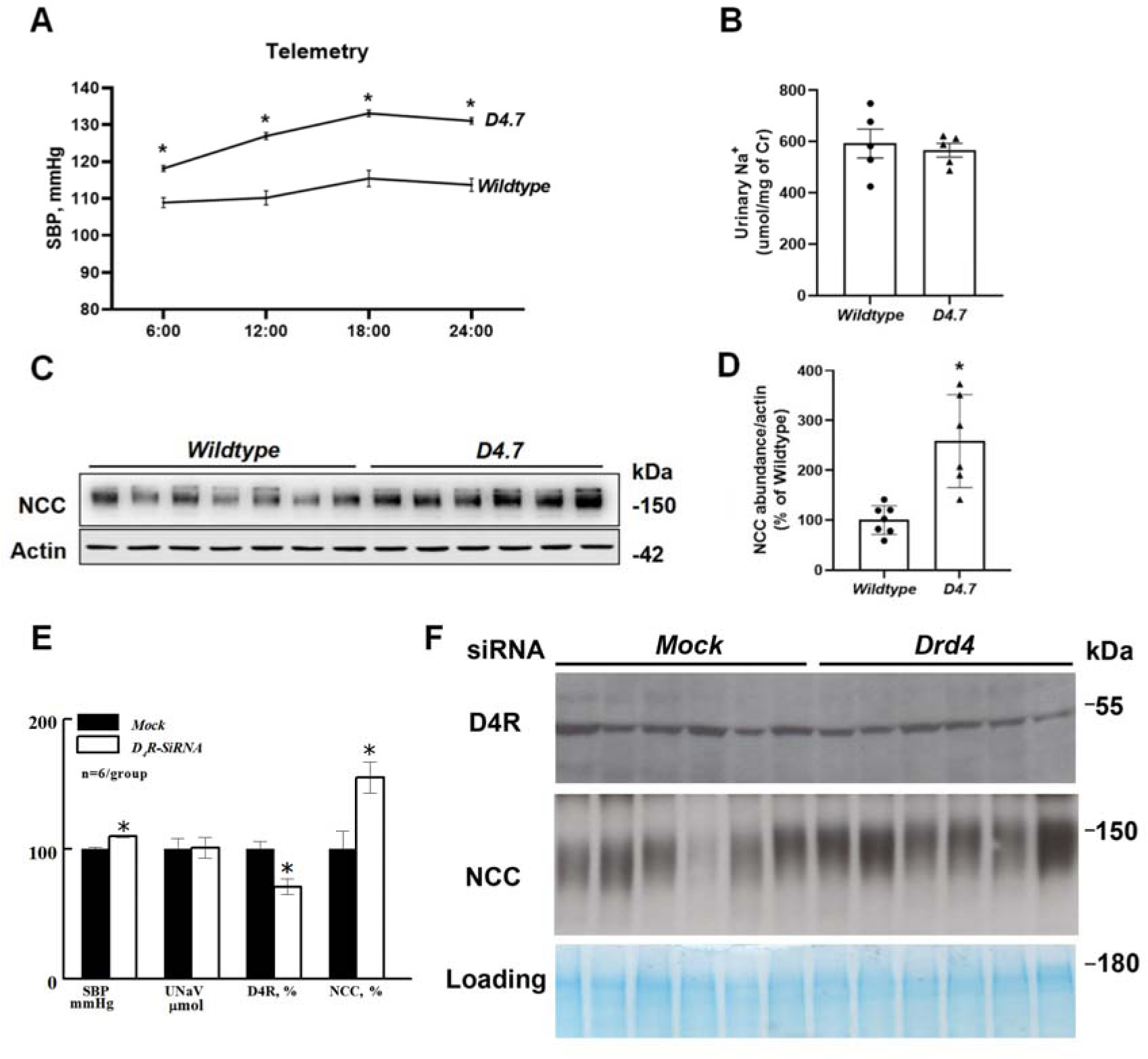
SBP, urinary sodium excretion, renal NCC protein abundance of *D4.7* knock-in mice or *Drd4*-siRNA treated mice. A. Systolic blood pressure (SBP), measured by telemetry, was modestly higher in *D4.7* knock-in mice (n=5) than in wildtype (n=5) mice. * *P* <0.05, *D4.7* vs. wildtype mice, Student’s *t*-test B. Urinary sodium excretion (UNaV) in *D4.7* and wildtype mice. There was no difference between the two groups. Student’s *t*-test between *D4.7* vs. wildtype mice. n = 5/group. C. Immunoblotting of NCC and actin in whole kidney homogenates from *D4.7* knock-in and wildtype mice. D. Quantitative analyses of the protein abundances observed in the immunoblots. The protein abundances of NCC were higher in *D4.7* knock-in and wildtype mice. \**P* < 0.05, vs wildtype, n = 6-7/group, Student’s *t*-test. E&F. SBP (mmHg, under anesthesia), UNaV (μmol/day) and renal D4R and NCC in kidney-specific-Drd4-siRNA treated mice. D4R was inhibited by 30% while BP and renal NCC increased in the mice without changes in urine sodium excretion. * *P* <0.05, n = 6/group, Student’s *t*-test.

## DISCUSSION

This study demonstrates that lack of D4R function, in *Drd4* deficient mice leads to aldosterone independent upregulation of the protein and activity of renal NCC that is reflected by increased HCTZ-sensitive natriuresis. A similar increase in total NCC protein abundance is also seen in mouse DCT cells treated with *Drd4-*siRNA and in mice expressing the *D4.7* variant or with reduction of renal D4R by sub-capsular siRNA infusion. On the contrary, treatment with a D4R agonist mediates a reduction in total NCC in these cells. Those findings indicate a downregulation of D4R on NCC, which is also confirmed by their colocalization and co-immunoprecipitation both in *vivo* and in *vitro*. The increase in total NCC protein is accompanied by decreased ubiquitination of NCC in mouse kidneys and mouse DCT cells with *Drd4* gene deficiency. In contrast total NCC decreased in mouse DCT cells treated with a D4R agonist these cells also showed increased NCC ubiquitination.

The decreased ubiquitination of NCC in mice and mouse DCT cells with *Drd4* gene deficiency may be related to increased USP48 - an enzyme that promotes the deubiquitination of NHE3 induced by D3R, another D2-like dopamine receptor(18, 59) - since D4R-mediated NCC ubiquitination is associated with alterations of USP48 in *vivo* and in *vitro*. The mechanism for the reduction of NCC protein and activity were further characterized and showed that D4R agonism promoted-NCC internalization, lysosome relocation and degradation.

The short-term inhibition of NCC by D4R may be caused by a decrease in the trafficking of NCC to the plasma membrane, similar to that proposed for WNK4(55). D4R agonism acutely decreases NCC signal in the plasma membrane resulting in its internalization Moreover, the reduction of its abundance at the plasma membrane blocked by a D4R antagonist. The colocalization between D4R and NCC is increased after 1 h of PD168077 treatment. D4R, similar to other dopamine receptors(60), exhibits constitutive activity that is shown in the current study since D4R antagonist L-745,870 not only blocks the agonist’s inhibition on NCC protein and activity in mouse DCT cells, but also increases NCC protein abundance by itself with extended treatment.

Ubiquitination triggers endocytosis and subsequent degradation of NCC(61). Prior reports have demonstrated that the E3 ubiquitin ligase Nedd4-2 is implicated in NCC ubiquitination(16, 17, 54). However, there were no changes on in the protein abundances of Nedd4-2 in our experiments. In turn we found that the protein abundance of USP48 increased in *Drd4* deficient mice and mouse DCTs. A small but significant decrease of USP48 is also found in mouse DCT cells treated with D4R agonist for 24 h, implicating an involvement of USP48 in the deubiquitination process of NCC. This suggests that the regulation of NCC by D4R might be attained through modulation of the equilibrium between ubiquitination and deubiquitination. A direct role in D4R-mediated ubiquitination of NCC and whether or not other E3 ubiquitin ligases are involved in NCC ubiquitination remain to be determined. Other ubiquitin specific proteases are also involved in the abnormalities of proteins’ poly-ubiquitination and mono-ubiquitination in some dopamine related disorders(59).

The decrease in NCC abundance caused by D4R stimulation may result from increased NCC degradation in lysosomes. We found that the D4R promotes the colocalization of NCC with the lysosome marker LAMP1 that is greater than the D4R-promoted colocalization of NCC with proteasomes, that is an organelle involved in the degradation of ubiquitinated proteins(42). Incubation with CQ prevents the D4R-mediated degradation of NCC. D4R induced-internalization and reduce protein level of NCC may support our results of reduced NCC activity. The co-immunoprecipration of ubiquitin with NCC from mouse kidneys and mouse DCT cells exhibits high molecular bands around 200 kDa, suggesting that D4R may cause NCC poly-ubiquitination(62). The poly-ubiquitination of NCC with band at 200 kDa is also seen in other studies(37). Big size proteins may be degraded in lysosome rather than in proteosome(63), as is suggested in our current study on D4R-promoted NCC degradation.

In summary, this work shows a novel mechanism: D4R inhibits NCC in renal DCT to maintain normal sodium homeostasis and blood pressure balance. Stimulation of D4R facilitates ubiquitination, internalization, promotes lysosomal localization and degradation of NCC. USP48 may be involved in the promoted ubiquitination of NCC by D4R.

### Novelty and Significance

#### What is known?

- The thiazide-sensitive sodium chloride cotransporter (NCC) is the major apical sodium transporter located in the mammalian renal distal convoluted tubule (DCT).
- The amount of sodium reabsorbed in the DCT through NCC plays an important role in the regulation of extracellular fluid volume and blood pressure.
- Dopamine and its receptors constitute a renal antihypertensive system in mammals. The disruption of *Drd4* in mice causes kidney-related hypertension.

What New Information Does This Article Contribute?

- Data from the current studies determined an inhibition of D4R on NCC protein abundances and activities in cultured mouse DCT cells via siRNA silencing and D4R agonism/antagonism and in mice with *Drd4* gene knockout.
- The downregulation of NCC by D4R is caused by increasing NCC ubiquitination via inhibiting ubiquitin-specific-protease-48 and subsequent internalization, relocation into lysosomes and degradation.
- The mice expressing the human *D4.7* variant develops high blood pressure with enhanced NCC protein abundances in kidney.

Our study identified the D4R-related molecular regulators of NCC in renal distal convoluted tubules and a novel mechanism of NCC ubiquitination and degradation and the upregulation of D4R deficiency on NCC protein abundances and activities in mice with Drd4 knockout and expressing the human *D4.7* variant, thus finally the major effect of D4R on renal sodium reabsorption and blood pressure.

## FUNDING

XW and her colleagues are supported by National Natural Science Foundation of China (81970605). MZ is supported by Postgraduate Research & Practice Innovation Program of Jiangsu Province (JX10413970). ML, ZR and PE are supported Nanjing Medical Science and Technology Development Fund (YKK22254, YKK22251, YKK20218). KKU, LA, IA and XW have been supported by NIH grants: DK134574 and DK119652.

## AUTHOR CONTRIBUTIONS

MZ completed most in *vivo* experiments on mice and NCC activity/degradation on cells, analyzed and figured the data and drafted the manuscript. ML performed partial in *vivo* experiments on mice and confocal microscopy on kidney sections and cell slips, and analyzed, figured and drafted the parts. ZR completed the transition and maintenance of the 2 gene mutant mice in China, the in *vivo* and in *vitro* experiments of *D4.7* mice and purification and labeling of D4R and NCC antibodies, performed QPCR of the kidneys and cells, and analyzed, figured and drafted the parts. Based on their contributions to the manuscript, they share the first authorship equally.

WW performed experiments on co-immunoprecipitation, confocal microscopy, protein degradation and related data analyses. KKU measured NCC activity, internalization and lysosome colocalization in vitro and wrote part of the draft. LA performed telemetry and partial mouse study. IA revised the manuscript critically. YJ, PW, and YX screened *Drd4*-siRNA, purified NCC antibody, assisted with the animal work and performed H&E staining on mouse kidney sections. XW conceptualized, designed the entire study, performed part of cell work, wrote/revised the manuscript critically and is responsible for the integrity of all data included. All authors read and approved the final version of submitted manuscript.

## ACKNOWLEDGMENTS

The authors thank Dr. Pedro Jose for gifting *Drd4* knockout mice, Dr. Sergi Ferré for gifting *D4.7* knock-in mice, Dr. Peter Freidman for gifting the mouse DCT cells and Dr. Mark Knepper for gifting initial rabbit NCC antibodies and origin of NCC null mouse samples, reviewing an early draft of the manuscript and providing helpful advice. The authors also thank Jianteng Xu for measuring electrolytes in urine and serum, Gangyi Zhu for assisting mouse handling and Figdraw (www.figdraw.com) for providing drawing tools and features.

## Nonstandard Abbreviations and Acronyms

αNKA: α sodium-potassium ATPase
AUC: Area under the curve
BP: blood pressure
CCK: cell counting kit
co-IP: coimmunoprecipitation
Ccr: creatinine clearance rate
CHX: cycloheximide
CQ: Chloroquine
Cr: creatinine
DCT: distal convoluted tubule
Drd: dopamine receptor gene
Drd4-/-: dopamine receptor gene 4 knockout mice
ENaC: epithelial sodium channels
FRET: fluorescence resonance energy transfer
HCTZ: hydrochlorothiazide
IP: immunoprecipitated
L: L-745,870
LAMP1: Lysosomal membrane protein 1
Met: metolazone
NCC: sodium chloride cotransporter
Nedd4-2: Neural-precursor-cell-expressed-developmentally down-regulated protein 4-like
NHE3: sodium hydrogen exchanger 3
NKCC2: Na+-K+-2Cl-cotransporter
PD: PD168077
PM: plasma membrane-enriched fractions
PSMC6: Proteasome 26S subunit, ATPase, 6
SBP: Systolic blood pressure
SEM: standard error of the mean
TCL: total cell lysates
USP48: ubiquitin-specific-peptidase
UNaV: Urinary sodium
Veh: vehicle
WKH: Whole kidney homogenates

## Notes

### Competing Interest Statement

The authors have declared no competing interest.

